# High-throughput retrieval of physical DNA for NGS-identifiable clones in phage display library

**DOI:** 10.1101/370809

**Authors:** Jinsung Noh, Okju Kim, Yushin Jung, Haejun Han, Jung-Eun Kim, Soohyun Kim, Sanghyub Lee, Jaeseong Park, Rae Hyuck Jung, Sang il Kim, Jaejun Park, Jerome Han, Hyunho Lee, Duck Kyun Yoo, Amos C. Lee, Euijin Kwon, Taehoon Ryu, Junho Chung, Sunghoon Kwon

**Author notes:** Equally contributed (Jinsung Noh, Okju Kim).

## Abstract

In antibody discovery, in-depth analysis of an antibody library and high-throughput retrieval of clones in the library are crucial to identifying and exploiting rare clones with different properties. However, existing methods have several technical limitations such as low process throughput from laborious cloning process and waste of the phenotypic screening capacity from unnecessary repetitive tests on the dominant clones. To overcome the limitations, we developed a new high-throughput platform for the identification and retrieval of clones in the library, TrueRepertoire^™^. TrueRepertoire^™^ provides highly accurate sequences of the clones with linkage information between heavy and light chains of the antibody fragment. Additionally, the physical DNA of clones can be retrieved in high throughput based on the sequence information. We validated the high accuracy of the sequences and demonstrated that there is no platform-specific bias. Moreover, the applicability of TrueRepertoire^™^ was demonstrated by a phage-displayed single-chain variable fragment (scFv) library targeting human hepatocyte growth factor (hHGF) protein.

## Introduction

*In vitro* display technologies enable fast selection of antibodies with desired properties (i.e., binding activity in most cases) from a highly diverse antibody library. The high diversity, which is derived from shuffling natural V_H_-V_L_ pairs or mutagenesis, enhances the possibility of finding potent clones in the libraries. Additionally, the association between the genotype and phenotype of clones in display systems allows for clone screening based on phenotype and, thereby, identification of the selected clones by genotype^1^. Among available *in vitro* display technologies, phage display is one of the most widely used techniques due to the high transformation capacity of phages and robustness of the process^2–4^. For example, the first FDA-approved fully human monoclonal antibody, adalimumab, was discovered through phage display^5^. Since then, many antibodies have been discovered and engineered through phage display, some of which have already been commercialized for clinical use^6^. In phage display-based antibody discovery, a phage display library, composed of diverse clones, is constructed. Then, the clones with binding activity to a target antigen in the library are enriched through a binding activity-based enrichment method, or biopanning. After several rounds of biopanning, the enriched clones are identified and retrieved for further characterization. In this step, identification and retrieval of large number of clones with enhanced “developability” are essential to increase the possibility to discover a prominent lead antibody that can be developed to a clinical drug. The term developability introduced in Jain, T. et al. include essential properties of a prominent lead antibody like high-level expression, high solubility, covalent integrity, conformational stability, colloidal stability, low poly-specificity, and immunogenicity of antibodies^7^.

However, in most cases, only a handful of major clones is identified since dominant clones usually outnumber rare clones and make the selected pool of clones relatively homogeneous. Clones with high developability can exist in rare population but can be lost due to technical limitations. Also, it was reported that clones with high binding activity to a target antigen can exist in a rare population in a library, due to the bias that results from the biopanning^8^. This was shown in our previous research, where certain rare clones in a library were retrieved and confirmed to have a binding activity to a target antigen^9^. Thus, to retrieve these rare clones with high developability and exploit them, in-depth analysis of a library is essential.

Improving throughput enables in-depth analysis of a library because sampling depth is increased. Conventionally, *in vitro* screening has been used to identify and retrieve clones in a phage display library^10^. Next-generation sequencing (NGS) technology was introduced to improve the screening process in terms of sampling depth^9,11,12^. However, the two existing methodologies, the *in vitro* screening and the NGS-based process, still have several technical limitations in identifying and retrieving rare clones in the library in high throughput. For *in vitro* screening, colony picking used for clone isolation is throughput-limited due to its labor intensiveness, unless a picking machine is used. Even when a picking machine is used, an additional cost for its operation is incurred, and a risk of cross-contamination still exists. Additionally, Sanger sequencing used in the genotyping step has throughput limitations derived from its laborious preparation step and high cost. Moreover, the process of genotyping after phenotyping causes the phenotype test to overlap with the same clones, which dramatically reduces the screening efficiency. Due to the low throughput and the limitations associated with the phenotypic screening capacity, only a handful of clones are identified and retrieved in most cases^3,13,14^.

To overcome these limitations, methods using NGS were introduced. Through high-throughput sequencing of a phage display library by NGS, high-resolution information about the composition of phage display libraries could be obtained. Based on the high-resolution information, rare clones in the libraries can be detected and confirmed to have binding activity toward the target antigens. Additionally, the change in the order of the process, from genotyping after phenotyping to phenotyping after genotyping, eliminated the loss of screening capacity in the phenotyping step. However, technical limitations associated with NGS itself still exist. First, the bias on the composition of clones in a library can be introduced during PCR amplification^15^. Second, NGS is vulnerable to sequencing and PCR errors^16^. Third, due to the short read length, linkage information between heavy and light chains of antibody fragments is lost^17^. Lastly and most importantly, to retrieve clones in the library for phenotypic testing, PCR rescue or *de novo* gene synthesis must be used. Low throughput of the methods makes the retrieval step costly and time-consuming, thereby limiting the number of clones to be tested in the following experiments^11,18^. Although unique molecular identifier (UMI) and single-cell sequencing techniques can be used to improve the NGS-based process, these techniques also rely on the low-throughput clone retrieval method^19–22^. In this respect, existing methodologies are not adequate for in-depth analysis of phage display libraries for detecting rare clones in and high-throughput retrieval of their physical DNA in the library for further characterization.

In this paper, we developed a new high-throughput cloning platform for antibody library screening named TrueRepertoire^™^ for in-depth analysis of phage display library and high-throughput retrieval of the clones’ physical DNA. Compared to conventional *in vitro* technologies, TrueRepertoire^™^ replaces the low-throughput colony picking system with a new high-throughput clone isolation system using a micro biochip and laser-based isolation method. Additionally, the low-throughput Sanger sequencing in the conventional process is replaced with a high-throughput clone identification system using multiplex PCR by barcoded primers, NGS, and barcode-based consensus sequence computation. Through TrueRepertoire^™^, highly accurate sequences of clones in an antibody display library are obtained while preserving linkage information between heavy and light chains of the clones. Additionally, a massive number of clones, including rare clones, in the library can be retrieved for use in the following experiments for further characterization (Fig. 1). To validate the performance of TrueRepertoire^™^, the accuracy of consensus sequences from the clone identification system of TrueRepertoire^™^ is demonstrated through cross-validation by Sanger sequencing. Then, we demonstrate, experimentally, that TrueRepertoire^™^ has no platform-specific bias derived from the new clone isolation system. In the experiment, an artificial library composed of clones with a pre-defined composition is used. Finally, the applicability of TrueRepertoire^™^ is demonstrated by applying TrueRepertoire^™^ to a phage-displayed scFv library.

**Fig.1.**
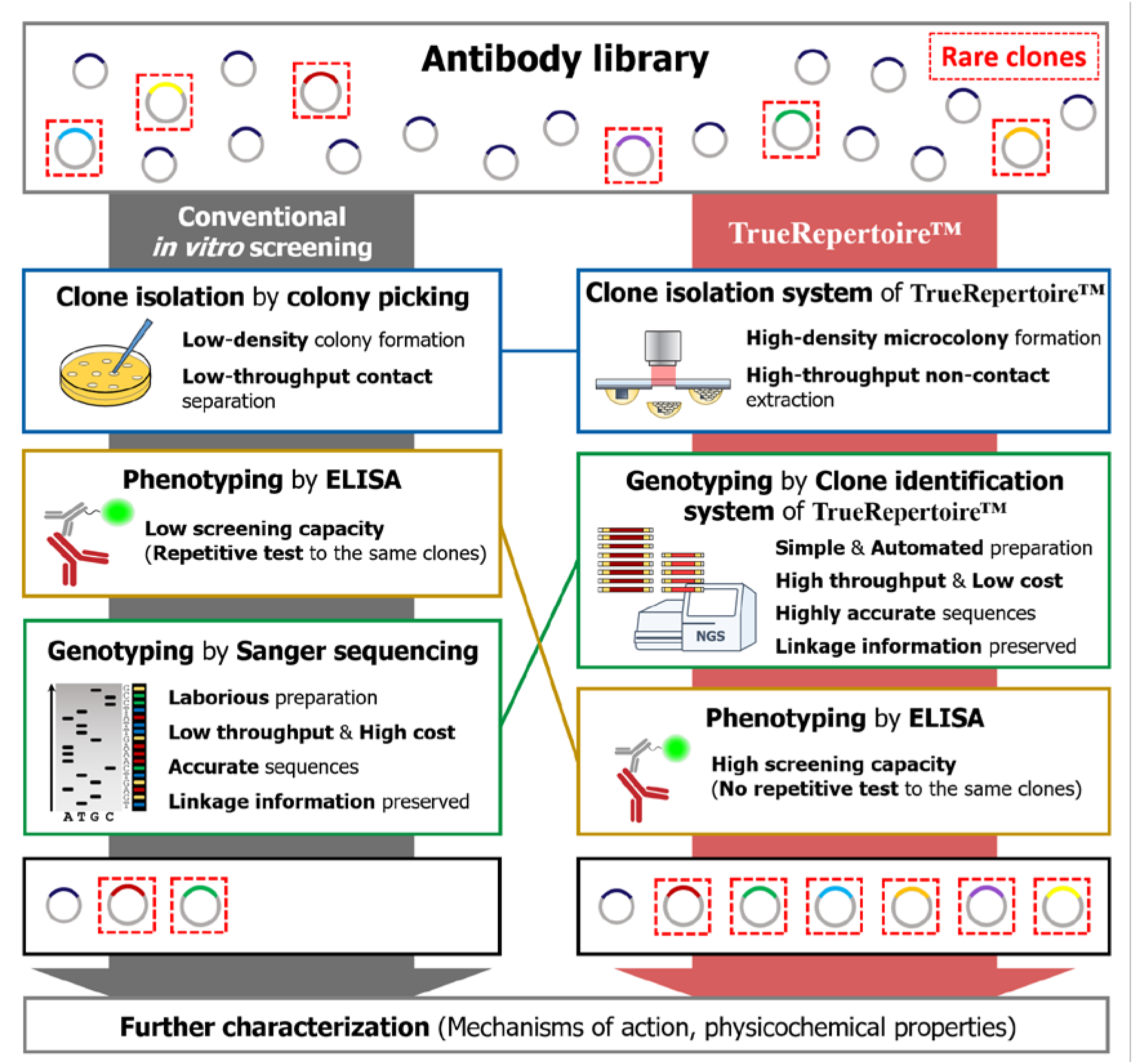
Comparison of the workflow between conventional *in vitro* screening and TrueRepertoire^™^. In the conventional *in vitro* screening process, clones in an antibody library are isolated by the colony picking method. The isolated clones are tested for their ability to bind to a target antigen by ELISA. Positive clones confirmed to have a binding activity are selected; then, the genotype of the clones is identified through Sanger sequencing. The low throughput and labor intensiveness of colony picking and Sanger sequencing limit the throughput. Moreover, the strategy of performing genotyping after phenotyping provokes the implementation of repetitive ELISA tests on the same clones, which reduces screening capacity. Overall, through the conventional *in vitro* screening process, in many cases, only a handful of clones are identified and retrieved for the following characterization. TrueRepertoire^™^ replaces colony picking with a new clone isolation system that enables the formation of microscale colonies in high density and extraction of the microcolonies in high throughput, without contact. The isolated clones are analyzed by the clone identification system of TrueRepertoire^™^. Through the clone identification system, highly accurate sequences of the clones are obtained based on the linkage information between heavy and light chains. Then, clones are selected based on the sequence information and tested for their binding activity by ELISA. Through the process, many clones, including rare clones, in the library can be identified and retrieved for further characterization.

## Results

### The micro biochip (SSICLE) and laser-based isolation of TrueRepertoire^™^

There are two important technological compartments that enable TrueRepertoire^™^: micro biochip and laser-based isolation. Conventional contact-based colony picking was replaced with a new laser-based colony isolation system. In the isolation system, a micro biochip, named as a Single-cell Separation and Incubation Chip capable of Light-mediated Extraction (SSICLE), is used to form intense amounts of microscale colonies on the chip. Following their formation, the microcolonies are extracted in high throughput from the chip using a pulsed infrared laser without contact.

SSICLE is composed of four main parts: an ITO-coated layer, a hydrophobic layer, hydrophilic pillar structures, and a cover glass. Microcolony-forming agarose droplets are formed around the pillar structures by self-assembly. Microorganisms are loaded into the agarose emulsion and grow to form microscale colonies. An aqueous solution consisting of the reagents for incubation, agarose for fixation, and microorganisms interacts with the hydrophilic pillar structures and the hydrophobic layer to form an agarose emulsion gathered around the pillar structures^23,24^. The emulsions are sealed with oil to prevent evaporation, and each emulsion acts as an incubator for the loaded microorganisms. The whole structures, including the agarose emulsions, are patterned on an ITO-coated glass cover. The ITO-coated layer of the chip reacts to an infrared laser and vaporizes to produce pressure for extracting the agarose emulsions (Fig. 2a).

**Fig.2.**
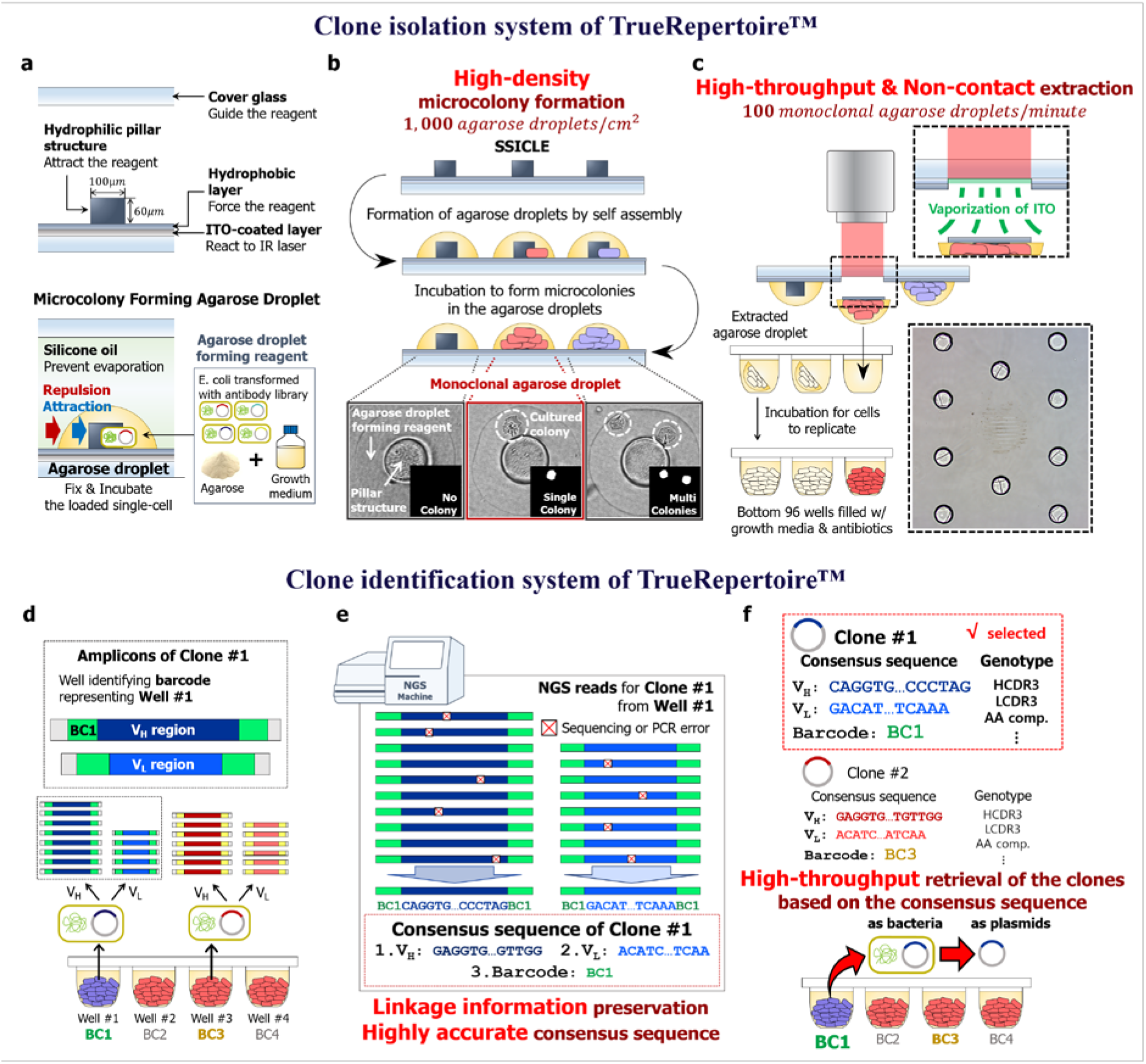
The clone isolation and clone identification systems of TrueRepertoire^™^. **a.** The structure of SSICLE. In SSICLE, microcolony-forming agarose droplets can be patterned in high throughput by self-assembly. The reagent comprising the agarose droplet interacts with the hydrophilic structure and hydrophobic layer to form agarose droplets. Single cells of transformants of the antibody library can be loaded into the agarose droplets and grow to form microscale colonies. **b.** Formation of the microcolonies on SSICLE. By using SSICLE, micro-colonies can be formed on the chip in high density. The monoclonal agarose droplets, each of which contains a single colony, are detected by an automated imaging and processing system. **c.** Extraction of the microcolonies by a laser. The ITO-coated layer of SSICLE reacts to an IR laser to extract the monoclonal agarose droplets. Through laser-based extraction, the microcolonies can be extracted in high throughput without contact. **d.** Multiplex PCR with barcoded primers of the isolated clones. Plasmids of the clones isolated in the previous step are amplified by multiplex PCR with barcoded primers. Through the amplification, amplicons with a pre-defined set of barcodes for each variable region are produced. **e.** Barcode-based sorting and computation of the consensus sequences. Based on the barcode information, NGS reads are sorted, and the consensus sequence corresponding to each clone is computed. In this step, sequencing and PCR errors can be excluded effectively. **f.** Genotype-based selection of the clones. Using the consensus sequence, clones are analyzed and selected based on the genotype of the clones. Then, the clones are retrieved by referring to the barcode information representing the position of the well containing the selected clone.

### The laser-based clone isolation system of TrueRepertoire^™^

As the first step of TrueRepertoire^™^, model organisms, especially bacteria, transformed with an antibody library are loaded onto SSICLE with adequate growth media for the model organisms and agarose for the fixation of cells. The chip loaded with the transformants is incubated to form micro-colonies in the agarose emulsions. After incubation, the number of colonies in each incubator is counted by an automated imaging and processing system to confirm the monoclonality of the agarose emulsion (Fig. 2b). Agarose emulsions with a single colony are detected in the previous step and are extracted from the chip into the bottom of 96 wells by a pulsed infrared laser (Fig. 2c). This non-contact-based extraction method using a pulsed laser eliminates the risk of cross-contamination. Additionally, the extraction step is fully automated, thus enabling high-throughput extraction of the clones (100 clones/minute, Supplementary Video. 1).

### The clone identification system of TrueRepertoire^™^

After incubation of the bottom wells containing the isolated colonies in the clone isolation step, plasmids of the isolated colonies are amplified by multiplex PCR for the amplification of variable regions with barcoded primers. The barcodes of the primers contain plate and well information, which enables barcode-based sorting of NGS reads (Supplementary Fig. 1). The PCR products are pooled and sequenced through the Illumina NGS platform (Fig. 2d). Reads from the NGS run are analyzed based on the barcode information of the reads. The NGS reads are aligned according to the barcodes; then, a consensus sequence representing each well is computed. In this step, a conservative threshold value for the minimum number of reads for computing the consensus sequences is used to prevent the effect of errors induced during PCR and NGS (Supplementary Fig. 2)^25,26^. The barcode-based amplification and following analysis provide several advantages over the conventional high-throughput sequencing of antibody libraries by NGS. First, the bias induced in the PCR amplification step is eliminated. Second, highly accurate sequences of clones are obtained. Third, linkage information between heavy and light chains of clones is preserved (Fig. 2e).

The consensus sequences are used in the following genotypic analysis of the clones. Based on the results of the genotypic analysis, clones to be further characterized are selected and then retrieved by referring to the barcode information of the selected clones. The clones can be retrieved in the form of bacteria or plasmids (Fig. 2f).

### Validation of the clone identification system of TrueRepertoire^™^

The clone identification system of TrueRepertoire^™^ involves multiplex PCR with barcoded primers, NGS, and barcode-based reads analysis for computing the consensus sequence of clones. Through the clone identification system, consensus sequences for clones are computed and used for further genotypic analysis of the clones.

To demonstrate the accuracy of the consensus sequences, the sequences were cross-validated by Sanger sequencing. First, clones in phage-displayed scFv libraries were isolated. Then, the isolated clones were sequenced through the clone identification system of TrueRepertoire^™^ to obtain consensus sequences of the clones. Additionally, plasmids of the clones were extracted and sequenced through Sanger sequencing. Then, the consensus sequences of TrueRepertoire^™^ were compared with V_H_ and V_L_ regions of the sequences, derived from Sanger sequencing for the cross-validation (Supplementary Fig. 3). A total 454 of consensus sequences of clones were cross-validated by Sanger sequencing via the abovementioned process. All 454 consensus sequences were perfectly matched to V_H_ and V_L_ regions of the corresponding Sanger results.

**Fig.3.**
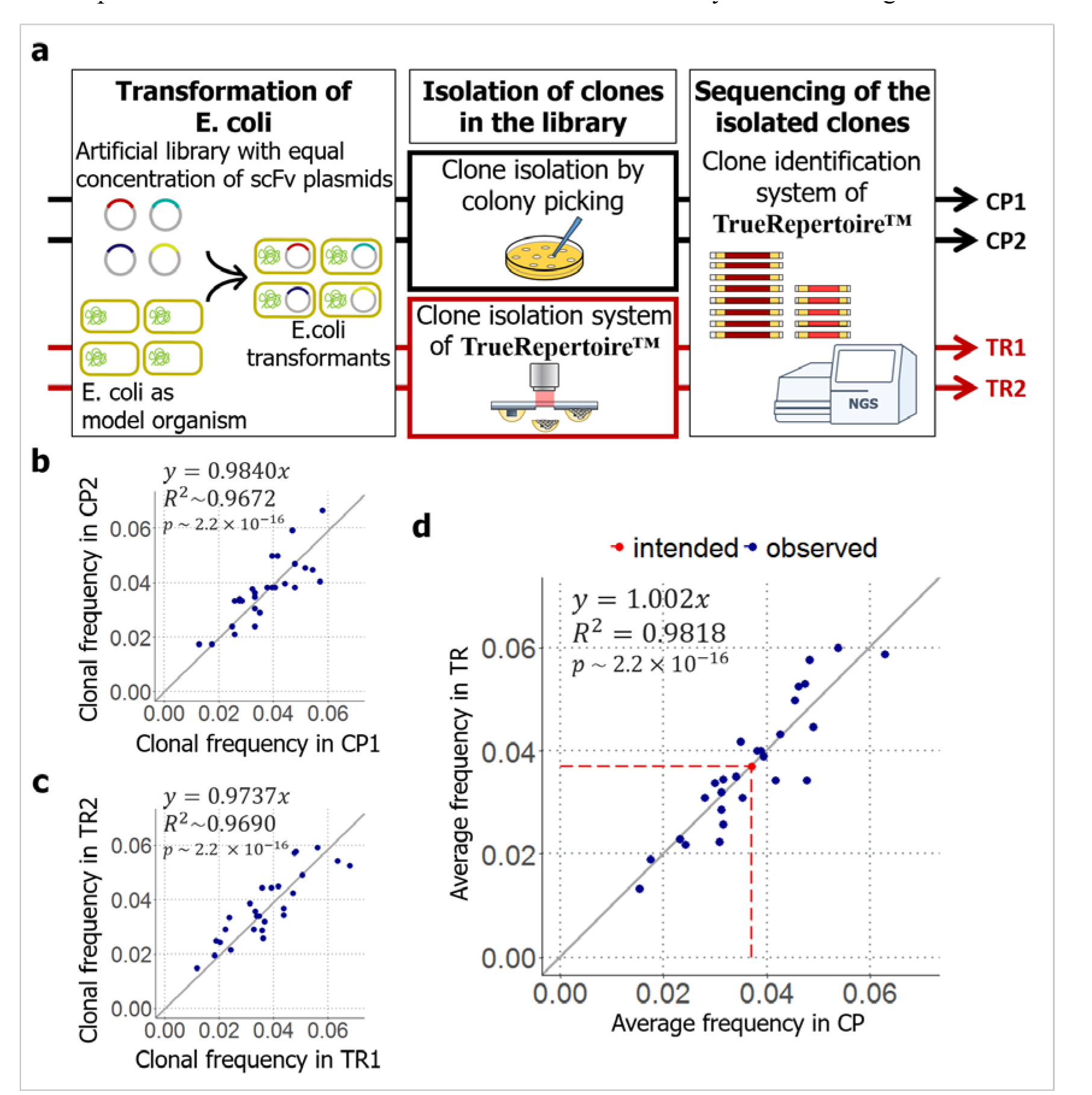
Validation of the clone isolation system of TrueRepertoire^™^. **a.** Workflow of the validation experiment. First, an artificial library with an equal concentration of scFv plasmids was transformed into *E. coli*. To determine whether the intended distribution of clones in the library is reproduced in TrueRepertoire^™^, the library was analyzed by the platform. As a control experiment, clones in the library were also isolated by colony picking. Then, the isolated clones were sequenced through the clone identification system of TrueRepertoire^™^. Two technical replicate experiments were conducted for each experimental condition. **b.** Reproducibility of colony picking. To determine how the results are reproduced in colony picking, the distributions acquired from the two replicate control experiments were compared. **c.** Reproducibility of TrueRepertoire^™^. To confirm the reproducibility of the platform, the results obtained from the two replicate experiments were compared. **d.** Comparison between the results of TrueRepertoire^™^ and the control results. To determine whether a platform-specific bias exists in TrueRepertoire^™^ derived from the new clone isolation, the averaged clonal frequencies of two replicates of the platform were compared with the averaged clonal frequencies of two replicates of the control using colony picking.

### Validation of the clone isolation system of TrueRepertoire ^™^

TrueRepertoire™ uses a new method for the isolation of clones in an antibody library. The new method comprises SSICLE and non-contact extraction of colonies by a laser. Through the new clone isolation system, clones in the library can be isolated in high throughput.

To confirm that there is no platform-specific bias derived from the new clone isolation system, an artificial library with a pre-defined composition of clones is constructed. The library is composed of an equal concentration of plasmids, each containing a DNA sequence for the expression of a scFv molecule. Individual clones of the artificial library are selected randomly from a naïve phage-displayed human scFv library (Supplementary File. 1). To determine whether the clonal frequencies, which are intended to be uniform, of the library are reproduced, the artificial library was transformed into the ER2738 *E. coli* strain by electroporation. TrueRepertoire^™^ was then applied to the library. As a control, the conventional colony picking was also applied to the same library in parallel. Clones in the library were isolated by colony picking and then analyzed by the clone identification part of TrueRepertoire^™^ to achieve high throughput. Two technical replicate experiments were conducted for each condition: an experiment using TrueRepertoire^™^ and a control experiment using colony picking.

For the control, 1,103 and 1,415 clones for each technical replicate were isolated by colony picking following a general protocol described previously^27^. Then, the isolated clones were analyzed by the clone identification system of TrueRepertoire^™^. The results indicated that the clonal frequencies of the clones were reproduced in the replicates (*p*-value=0.8622, chi-squared test between the replicates). However, the distribution slightly deviated from the intended uniform distribution (D_KL_ = 0.0441, KL-divergence between the distribution of clonal frequencies from colony picking and the uniform distribution of unif {1, 27}). Because the slight deviation was reproduced in both replicates, the deviation might have stemmed from factors inducing bias in the library composition other than a random effect. A possible factor contributing to the bias is thought to be the variation in transformation efficiencies with plasmid form (Supplementary Fig. 4)^28^.

For the experimental condition using TrueRepertoire^™^, 2,067 and 2,078 clones for each technical replicate were isolated and their genotypes were identified as described in the methods section. As a result, the clonal frequencies of the clones were reproduced in both replicates as in the control experiment (*p*-value=0.1069, chi-squared test between the replicates, Fig. 2c). The results obtained from the replicate experiment confirmed the reproducibility of TrueRepertoire^™^. A deviation from the uniform distribution was also observed (D_KL_ = 0.0560 KL-divergence between the distribution of clonal frequencies from TrueRepertoire™ and the uniform distribution of unif {1, 27}).

To confirm the existence of platform-specific bias, the clonal frequencies from TrueRepertoire^™^ were compared with the clonal frequencies from the control experiment using colony picking. To make the comparison, averaged clonal frequency values of the two replicates for each condition were used. The averaged clonal frequencies of the clones identified through the two platforms, TrueRepertoire^™^ and colony picking, were confirmed to follow the same distribution (p-value=0.9578 in chi-squared test between the averaged clonal frequencies of two replicates from colony picking and TrueRepertoire^™^, Fig. 2d). In addition, clonal frequencies from the individual experiment for each condition, TrueRepertoire^™^ and colony picking, were also compared (Supplementary Fig. 5).

All clones in the library were identified through TrueRepertoire^™^. A distortion of clonal frequencies from the intended frequencies, the uniform distribution of unif{1,27}, was observed among the results yielded by TrueRepertoire^™^. However, the distortion was also observed in the results of the control experiment using colony picking, and the degrees of distortion for the platforms were similar (D_KL_ = 0.0441 for colony picking and D_KL_ = 0.0560 for TrueRepertoire^™^). For both platforms, the distortion is thought to be introduced by the variation in transformation efficiencies with the forms of clonal plasmids. Thus, compared with conventional colony picking, TrueRepertoire^™^ has no platform-specific bias introduced by the new clone isolation system.

### Application of TrueRepertoire^™^

To validate the applicability of TrueRepertoire^™^, TrueRepertoire^™^ was applied to a phage-displayed scFv library targeting human hepatocyte growth factor (hHGF) protein (Supplementary File. 2). The library was constructed from bone marrow of white leghorn chickens; the chickens were immunized with hHGF. Then, the library was enriched through five rounds of biopanning. The number of biopanning rounds was determined based on the output titer during the biopanning (Supplementary Table. 1). Then, the library was analyzed by TrueRepertoire^™^. Additionally, for the comparison, the library was analyzed by NGS. Each variable region of the clones, the V_H_ and V_L_ regions, was amplified separately by PCR. The amplicons were then sequenced through the Illumina Miseq platform.

A total of 3,377 clones were isolated and analyzed by TrueRepertoire^™^. Among the analyzed clones, there exist 955 unique clones at the amino acid sequence level. Without considering the linkage information between heavy and light chains, the results feature 325 unique V_H_ amino acid sequences and 905 unique V_L_ amino acid sequences. To confirm that the results acquired from TrueRepertoire^™^ reflect the real distribution of clones in the library, the results of TrueRepertoire^™^ were compared with those of NGS. However, in NGS, linkage information between heavy and light chains is lost. Thus, in the TrueRepertoire^™^ results, frequencies of V_H_ sequences were calculated by summing the clonal frequencies of the clones with the same V_H_ sequence and then compared with the results of NGS for the V_H_ region. The comparison confirmed that the clonal frequencies of clones from TrueRepertoire^™^ correspond with those from NGS (Fig. 4a). Additionally, it was analyzed whether linkage information acquired from TrueRepertoire^™^ was preserved in the NGS results. The linkages between heavy and light chains were analyzed for the 10 most frequent clones among the TrueRepertoire^™^ results. Among the 10 most frequent clones, only two clones, the most frequent and fifth frequent clones, maintained their linkage information between heavy and light chains (Table 1).

**Fig.4.**
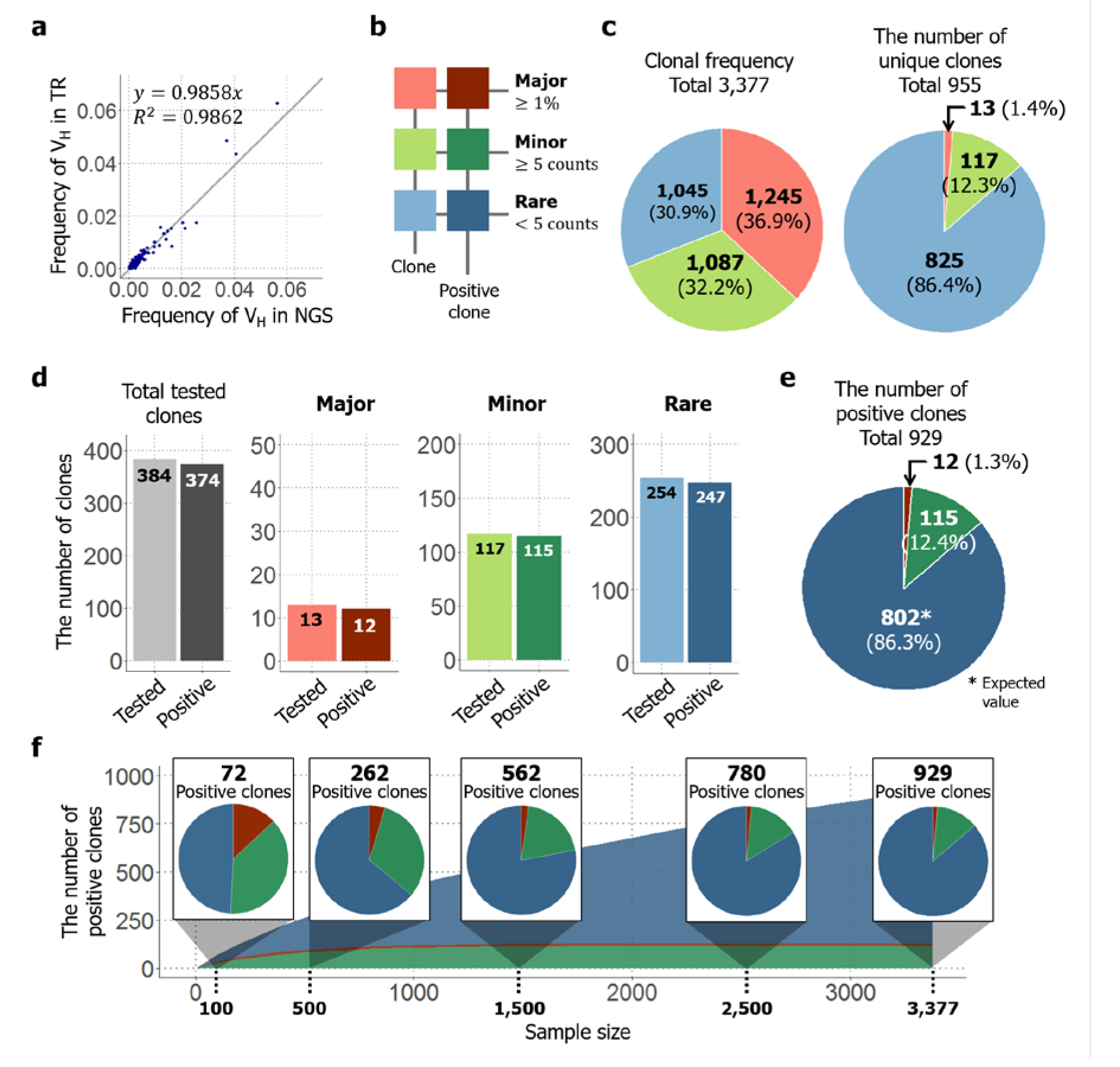
Application of TrueRepertoire^™^ to a phage-displayed scFv library targeting hHGF protein. **a.** Comparison of the frequencies of a heavy-chain sequence of the identified clones between the NGS and TrueRepertoire^™^ results. To confirm that the clonal frequencies of the clones identified through TrueRepertoire^™^ reflect the real distribution of the library, the frequencies of the heavy-chain sequence were calculated and compared with those from NGS. **b.** Groups of the identified clones. To clarify the distribution of the library, the clones were categorized into three groups, major, minor, and rare, according to the clonal frequency of the clones. The major group was denoted in red, the minor group in green, and the rare group in blue. Positive clones having binding activity to hHGF of each group were denoted in darker colors. **c.** Composition of the clonal frequencies of the groups and the number of unique clones in each group. The clonal frequency of the groups refers to the sum of clonal frequencies of the clones in each group. **d.** Binding activity test of the clones by ELISA. A total of 384 clones were retrieved and then tested for their binding activity to hHGF protein. All clones in the major and minor groups were retrieved and then tested for their binding activity. In the rare group, 254 clones were selected randomly from the total number of clones in the group and then tested for their binding activity. Among the tested 384 clones, 374 clones were confirmed to be positive clones with binding activity. **e.** The expected composition of total positive clones in the library. The expected number of positive clones in the rare group was estimated based on the positive rate from the previous ELISA test. **f.** The results of the sampling simulation. To determine how the number of positive clones and the composition of the positive clones changes, as the throughput of TrueRepertoire^™^ increases, a sampling simulation was conducted.

**Table 1.** Top 10 clones with respect to clonal frequency yielded by TrueRepertoire^™^ Rank of the clones was calculated based on clonal frequencies in the TruRepertoire^™^ result. In the V_H_ - V_L_ pair column, “H” and “L” denote heavy and light chains each. The numbers following “H” and “L” denote the rank of the respective chains with the respect of the corresponding read frequencies in the NGS results. The result of the binding activity test using phage ELISA was also represented.

| TrueRepertoire™ |  |  | NGS |  | ELISA |
| --- | --- | --- | --- | --- | --- |
| Rank | V<sub>H</sub>-V<sub>L</sub> pair | Clonal frequency (%) | V<sub>H</sub> frequency (%) | V<sub>L</sub> frequency (%) | Binding activity (O/X) |
| 1 | H1-L1 | 9.27 | 13.08 | 3.20 | O |
| 2 | H6-L8 | 4.83 | 3.56 | 0.96 | O |
| 3 | H2-L5 | 3.67 | 12.67 | 1.17 | O |
| 4 | H3-L13 | 2.37 | 7.62 | 0.65 | O |
| 5 | H9-L9 | 1.63 | 1.95 | 0.83 | O |
| 6 | H1-L40 | 1.54 | 13.08 | 0.31 | O |
| 7 | H14-L92 | 1.27 | 1.26 | 0.18 | O |
| 8 | H7-L47 | 1.18 | 2.45 | 0.28 | O |
| 9 | H2-L30 | 1.13 | 12.67 | 0.40 | O |
| 10 | H12-L31 | 1.10 | 1.39 | 0.37 | O |

To clarify the composition of the library, the identified clones were categorized into three groups— major, minor, and rare—according to the clonal frequency of the clones. Clones with a frequency greater than 1% of the total number of analyzed clones were categorized into the major group. For the remaining clones, if a clone had a clonal frequency of more than or equal to five, the clone was categorized into the minor group, and if the frequency was less than five, the clone was categorized into the rare group (Fig. 4b).

As previously mentioned, 955 unique clones existed among the 3,377 analyzed clones. The clonal frequencies of the groups were calculated by summing the clonal frequencies of the clones in each group, and the values were confirmed to be similar to each other. Although the different groups showed approximately equal values with respect to clonal frequency, the number of unique clones in the rare group represented 86.4% of the total number of unique clones identified (Fig. 3c). To calculate the number of positive clones with binding activity toward hHGF in each group, clones were selected from each group, then tested for their binding activity by phage ELISA. In total, 384 clones were tested as described in the methods section, among which 374 clones were confirmed to have binding activity toward hHGF (Supplementary Fig. 6). All 13 clones in the major group and all 117 clones in the minor group were tested; 12 clones in the major group and 115 clones in the minor group were confirmed to have positive signals in the test. Regarding the rare group, 254 clones were selected randomly from among those in the rare group and then tested for their binding activity. Among the 254 clones, 247 clones were confirmed to have a binding activity (Fig. 4d). Using the positive rate from the test for the rare group, the expected number of positive clones in the rare group was estimated to be 825. Clones not only from the major and minor groups, but also from the rare group had a high positive rate in terms of binding activity. Thus, the rare group was also dominant in terms of the number of positive clones, like the case of the number of unique clones (Fig. 4e). Additionally, to determine how the number of positive clones and the composition of the positive clones change as the throughput of TrueRepertoire^™^ increases, a sampling simulation was conducted. Data points were sampled randomly without replacement from the whole. Then, the expected number of positive clones in the sample was calculated based on the results of the phage ELISA test. Sampling was conducted in every 100 sample size from 100 to the whole sample size, 3,377. Sampling was performed 1,000 times at each sample size, and averaged values were then calculated and represented. As the through-put of TrueRepertoire^™^ increased, the number of positive clones increased following an approximately logarithmic scale. For the composition of the positive clones, at a sample size of 100, the number of positive clones from the major and minor groups represented more than 50% of the total number of positive clones. As the sample size increased, while the number of positive clones from the major and minor groups saturated early in the simulation, the number of positive clones from the rare group increased continuously. Eventually, clones from the rare group dominated among the positive clones in the latter phase of the simulation (Fig. 4f).

To assess the functional diversity of the clones, heavy-chain complementarity determining region 3 (HCDR3) of the clones was extracted and analyzed. A total of 115 unique HCDR3 clusters was identified, and a phylogenetic tree of the region was drawn following the analytical method described previously^29,30^. The clusters had unique features in the use of amino acids and the length of the HCDR3 region (Fig. 5a).

Additionally, the clonal frequency and number of unique clones corresponding to each HCDR2 cluster were represented together (Fig. 5b, c). The clonal frequency of the clusters is the sum of clonal frequencies of clones having the HCDR3 amino acid sequence of the cluster. Considering the mechanism of biopanning, HCDR3 sequences of clusters with a high clonal frequency are expected to dominate through the biopanning process, possibly due to its superior affinity or higher stability. The clonal frequency can be used for prioritizing clones to be retrieved and further characterized. The number of unique clones in a cluster refers to the number of unique clones having the HCDR3 amino acid sequence of the cluster. This value can also be interpreted as the number of different retrievable clones having the HDCR3 sequence of the cluster. After capturing the possible number of retrievable clones of each cluster based on this value, the genotype of the clones in the group, rather than the HCDR3 sequence alone, can be analyzed. Then, the results of the genotype-based analysis can be used for prioritizing the clones based on the presence of post-translational modification motif, isoelectric point, or the expected immunogenicity.

**Fig.5.**
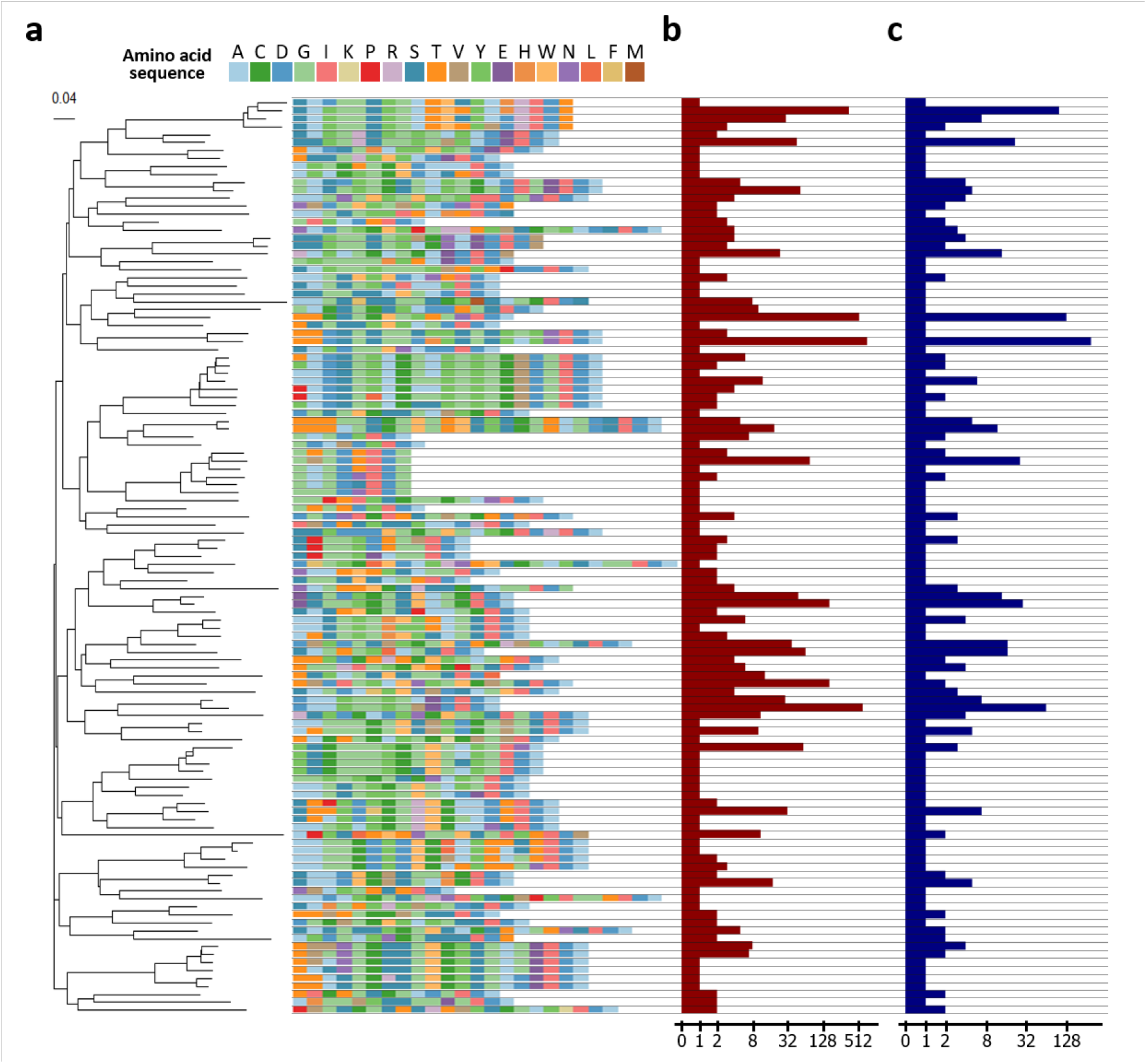
Analysis of HCDR3 region of the clones. **a.** HCDR3 clusters of the clones and a phylogenetic tree of the clusters. To interrogate diversity at the functional level, the clones were clustered based on the sequences of the HCDR3 region. A total 115 unique clusters were identified and confirmed to have a diverse composition of amino acid sequences of the HCDR3 region and a heterogeneous length of the HCDR3 region. Then, the tree was built by ClustalW2 using the neighbor joining method. The scale bar indicates the probability of substitution. **b.** Clonal frequencies of the clusters. Clonal frequencies of the cluster represent the sum of clonal frequencies of the clones having the HCDR3 sequence of the clusters. The clonal frequency can be interpreted as the importance of a cluster with respect to binding activity. In the following genotype-based screening step, this value can be used. **c.** The number of unique clones in the cluster. This value represents the number of unique clones having the HCDR3 sequence of the cluster. Based on this value, the number of different clones with a specific HCDR3 sequence can be captured. The clones can be screened based on the genotype of the clones.

## Discussion

In-depth analysis of a phage display library and retrieval of clones in high throughput are crucial in identifying and exploiting prominent clones with different characteristics such as mechanism of action or physicochemical properties of antibody fragments. Although clones with binding activity are enriched through biopanning, there exist other characteristics of antibodies that determine the clinical use^31^. Those include high-level expression, high solubility, covalent integrity, conformational stability, colloidal stability, low polyspecificity, and immunogenicity of antibodies, which were termed “developability” in previous research^7^. Thus, clones with desired characteristics as a drug that are not enriched enough during biopanning, can exist as rare clones in a phage display after several rounds of biopanning. To identify and exploit these prominent rare clones, in-depth analysis of a phage display library and high-throughput retrieval of clones should be followed after biopanning of the library.

Here, we introduced a new robust and high-throughput platform, TrueRepertoire^™^, for the identification and retrieval of clones in a phage display library. TrueRepertoire^™^ is composed of two systems: clone isolation system and clone identification system. In the clone isolation system, microcolonies of clones in the library can be formed in high density using the micro biochip SSICLE, and the microcolonies can be extracted from the chip in high throughput without contact by laser. The isolated colonies can be analyzed by the clone identification system of the platform to compute highly accurate consensus sequence of the clones while preserving the linkage information between heavy and light chains of the clone. By using the novel systems, TrueRepertoire^™^ can identify and retrieve a massive number of clones, including rare clones, from a phage display library within a short period and at a relatively low cost compared to the existing technologies. We identified and retrieved 955 unique clones from the phage-displayed scFv library targeting hHGF by using TrueRepertoire^™^. To retrieve the same number of clones through NGS followed by *de novo* gene synthesis, it costs more than 10-fold compared to the cost of TrueRepertoire^™^. In addition, as shown in the application of TrueRepertoire^™^ section, while the linkage information between heavy and light chains is preserved in TrueRepertoire^™^, the linkage information is lost in NGS. Thus, TrueRepertoire^™^ saves significant effort and time by preserving the real linkage between heavy and light chains of clones. Through this paper, it was validated that the consensus sequences have high accuracy and there is no platform specific bias in TrueRepertoire^™^. Also, we succeeded in identifying and retrieving many rare clones by applying the platform to the phage-displayed scFv library targeting hHGF. The retrieved clones were confirmed to have binding activity to the target antigen, hHGF. In addition, the functional diversity of the identified clones in the library was shown by analyzing HCDR3 region of the clones. The clones had heterogeneous HCDR3 in the aspect of sequence length and amino acid composition. Considering the heterogeneity, the clones are expected to bind to different epitopes of hHGF, as reported before^32^.

TrueRepertoire^™^ enables high-throughput genotype-based screening of the clones in a phage display library; thus clones to be retrieved for further characterization can be prioritized, which reduces the cost for the following characterization of the clones. In conventional antibody drug discoveries, early screening and optimization steps focus on characteristics such as binding activity, potency, and stability for selection of lead constructs, while pharmacokinetic properties which can influence both efficacy and toxicity are typically characterized later in development and on a small number of lead monoclonal antibody constructs. Predicting pharmacokinetic properties of antibodies in the genotype-based screening can reduce the time needed for drug discovery and development by improving the lead monoclonal antibody selection process^33^. TrueRepertoire^™^ provides a highly accurate consensus sequence of the clones. Additionally, the clonal frequency of the clones can be calculated based on the consensus sequences without a platform-specific bias. Using the consensus sequence and clonal frequency of the clones, characteristics of clones, including pharmacokinetics of antibodies, can be expected. Therefore, clones can be selected based on genotype-based screening results, and the selected clones can be retrieved by referring to the barcode information of the consensus sequence^7,33,34^. As a representative example of the genotype-based screening through TrueRepertoire^™^, charge distribution of antibodies can be analyzed by the consensus sequence of the clones, then clones expected to have poor pharmacokinetics can be excluded in the retrieval. There have been reports of antibodies with a high positive charge having poor pharmacokinetic profiles, and removing or repositioning the positive charges or counterbalancing with negative charges dramatically improved the pharmacokinetic profiles^35–37^. The results of those reports can be introduced in the genotype-based screening to prevent clones with poor pharmacokinetics from being further characterized in the following development. Thus, the efficiency of the development process can be enhanced.

Moreover, TrueRepertoire^™^ can enhance the screening efficiency of ELISA which is followed for confirmation of binding activity of the retrieved clones. In the conventional *in vitro* screening, identification of the clones through Sanger sequencing is conducted after a phenotypic screening of the clones by ELISA, which provoke repetitive tests to the same clones. The waste of screening capacity increases the cost of the whole process and can limit the number of clones to be further characterized in the development. TrueRepertoire^™^ can provide refined sets of clones to be tested without overlap through sequence confirmation and genotype-based screening by using the consensus sequence of the clones, which can prevent repetitive tests and increase the screening capacity of the process. When the recombinant expression of a target antigen is not available, the high screening capacity of the platform acts as a critical advantage.

TrueRepertoire^™^ can also be applied to libraries other than a phage display library. The platform can be used to analyze such libraries and retrieve clones in those libraries. If a library is in a plasmid form or other forms that can be transformed into a model organism (i.e., bacteria), clones in the library can be identified and retrieved through TrueRepertoire^™^. For example, antibody libraries used in other than phage display systems or protein libraries used in the directed evolution of a protein can be processed with TrueRepertoire^™^. Additionally, TrueRepertoire^™^ can be used in microbiome research, particularly to identify and retrieve bacteria existing in the environment.

In summary, we developed a new robust platform, TrueRepertoire^™^, to identify and retrieve clones in high throughput in a phage display library. It was shown that there is no platform-specific bias in TrueRepertoire^™^, and the applicability of the platform was demonstrated by finding many effective rare clones in a phage-displayed scFv library. TrueRepertoire^™^ provides a platform for high-throughput genotype-based screening of clones, which provides several advantages over previously developed processes. Additionally, TrueRepertoire^™^ can be applied to antibody libraries other than phage display libraries; protein libraries, which are used in the directed evolution of a protein; and microbiota native to a given environment.

## Methods

### Fabrication of the Single-cell Separation & Incubation Chip capable of Light-mediated Extraction (SSICLE)

To remove a contaminant on the surface, an indium tin oxide (ITO) deposited glass is washed thoroughly with absolute ethanol. To make a hydrophobic layer, 5% v/v solution of photoinitiator (2-Hydroxy-2-methylpropiophenone, Sigma Aldrich) and PEGDA250 (Polyethylene glycol diacrylate, M_n_ 250, Sigma Aldrich) is spin-coated on the ITO deposited glass (500 rpm – 5 s, 3000 rpm – 30 s, A/Del time – 5s). After the coated hydrophobic layer is cured by ultraviolet light (365 nm, 85 mW, 10 s), residual uncured monomers of PEGDA250 were washed with absolute ethanol. To make 60 um of pillar structures, the same height of spacers were attached on both ends of the hydrophobic layer coated glass, then a photomask for the pillar structure was put on the spacers. Then, 5% v/v solution of the photoinitiator and PEGDA700 (Polyethylene glycol diacrylate, M_n_ 700, Sigma Aldrich) was loaded to the gap between the photomask and the hydrophobic layer coated glass. The pillar structures (60um height) were patterned on the hydrophobic layer coated glass by irradiating ultraviolet light on the photomask (365 nm, 85 mW, 5 s), then residual uncured monomers were washed with absolute ethanol. After making the spacers (180um height) on both ends of the pillar structure patterned glass, a slide glass was put on the spacers and fixed to guide a solution loaded to the chip.

#### The clone isolation system of TrueRepertoire^™^

##### Transformation *E. coli* with an antibody library

To make transformants of an antibody library, E.coli (ER2738, Lucigen) was transformed with 1 ng of plasmid DNA for the library by electroporation according to the protocol from the manufacturer of the electroporation apparatus (MicroPulser^™^ Electroporator, Bio-rad).

##### Formation of the microcolony forming agarose droplets in SSICLE

The transformants were diluted with 1% m/m low gelling temperature agarose SOC solution with antibiotics. The diluted transformants solution was loaded into SSICLE, then washed with silicone oil (Sigma Aldrich). Through this two-step manipulation, the microcolony forming agarose droplets were formed in high-throughput in SSICLE by self-assembly. After cooling the agarose droplets to fix the cells, the chip was incubated at 37°C overnight in a cell incubator.

##### Image processing of the chip

After the incubation, images for the whole area of the chip were taken with an automated imaging system. The images were converted to gray-scale and contrast of the images was enhanced for the following image processing. Microcolonies in the agarose droplets were detected through several rounds of morphological operations including erosion and dilation. After counting the number of microcolonies in each agarose droplet, agarose droplets with one microcolony were categorized as monoclonal agarose droplets, and the coordinates of the monoclonal agarose droplets were recorded for the following extraction step.

##### Extraction of the monoclonal agarose droplets

Using the coordinate information obtained in the image processing step, the monoclonal agarose droplets were extracted from the chip into 96-well filled with growth media and antibiotics by a fully automated in-house laser extraction system. The extraction system is comprised of stage controllers and a pulsed infra-red laser instrument. After the extraction, the bottom 96-well plates containing the extracted monoclonal agarose droplets were incubated overnight for *E. coli* in the agarose droplets to replicate enough.

#### The clone identification system of TrueRepertoire^™^

##### Multiplex PCR with barcoded primers and NGS

After the overnight incubation, variable regions of the plasmids of *E. coli* in the 96-well plates were amplified by multiplex PCR with two sets of primers. The primers contain barcode regions containing plate- and well-information of the amplified *E. coli*. The multiplex PCR (95 °C for 9 min 30 s, 26 cycles of 95 °C for 25 s, 60 °C for 25 s, and 72 °C for 1 min, 72 °C for 3 min) was performed with PCR reagent (10 μL of BioFACT^TM^ 2× Taq polymerase Master Mix, 5 μL of nuclease free water, 1 μL of 10 μM forward barcoded primer, 1 μL of 10 μM reverse barcoded primer, and 1 μL of the growth media containing *E. coli*). After the multiplex PCR, the PCR products were pooled and purified using magnet beads. Then, the purified PCR products were prepared for NGS run (Miseq, 2×300 bp, Illumina) following the protocol from the manufacturer.

##### Pre-processing and demultiplexing

Paired-end reads were merged using PEAR (Paired-End read merger, v0.9.6) and unmerged paired-end reads were eliminated^38^. For quality control, merged reads with a low median quality PHRED score were additionally filtered out. Remaining reads were demultiplexed by the barcode region sequence. Then, the reads were tagged with the type of chains (heavy or light) and well location corresponding to the barcode sequence.

##### Sequence clustering

For each group of reads tagged with the same chain type and well location, the reads were clustered through the following steps. First, primer sequence was trimmed for each read, and sequence abundances (read count) were calculated based on the occurrence of the remaining immunoglobulin sequences. Sequence clusters were constructed using a greedy algorithm that sets the most abundant sequence as the representative sequence for each cluster and assigns new sequences to the closest cluster based on edit distance if the distance is smaller than a threshold and otherwise allocates a new cluster.

##### Well filtering

After the sequence clustering, some of the well data were discarded if either the V_H_ or V_L_ group was under one of the following conditions. i) The relative abundance of the most abundant cluster is less than a threshold value. ii) The read counts of the most abundant cluster of a group is less than a threshold value which is computed by a statistical analysis. (Supplementary Figure. 2)

##### Extracting consensus sequence

Finally, a consensus sequence was constructed for each remaining group via the following steps. First, sequences in the most abundant cluster were aligned using Clustal Omega^39^. Next, the most abundant nucleotide for each base position was calculated and assembled together to make a consensus sequence. Gaps were ignored after the construction.

Every bioinformatics process in this section not designating a software name in the text was processed using an in-house software of MSSIC platform.

##### Immunization of chicken for generation of anti-hHGF antibody

Six white leghorn chickens were immunized and boosted twice with 5 μg of hHGF (100-39H, Peprotech, Rocky Hill, NJ, USA). Bloods were collected from the wing veins prior to immunization and during the immunization for ELISA to evaluate the immunization status.

##### Construction of combinatorial phage-displayed scFv library and biopanning

The chickens were sacrificed a week after the second boosting. Bone marrow was harvested for total RNA isolation. Using the RNA, cDNA was synthesized to generate a phage-displayed scFv library as described previously^40^. After the library construction, five rounds of bio-panning were performed against hHGF (PGA104, PanGen Biotech Inc., Suwon, Republic of Korea) conjugated magnetic beads as described previously^41^.

##### Phage ELISA

Clones obtained through TrueRepertoire^™^ were rescued by infection of helper phage and subjected to phage ELISA using hHGF coated microtiter plates (3690, Corning, NY, USA) to select binders as described previously^10^. Briefly, the microtiter plates were coated overnight at 4°C with 100 ng of hHGF (100-39H, Peprotech, Rocky Hill, NJ, USA) in coating buffer (0.1 M sodium bicarbonate, pH 8.6). The wells were blocked with 150 μL of 3% (w/v) bovine serum albumin (BSA) in phosphate-buffered saline (PBS) for 1 h at 37°C. The plates were then sequentially incubated with scFv-displaying phages in the culture supernatant and horseradish peroxidase (HRP)-conjugated mouse anti-M13 monoclonal antibody (GE Healthcare, Pittsburg, PA, USA) with intermittent washing using 0.05% (v/v) Tween 20 in PBS (PBST). Finally, the plates were washed again with 0.05% PBST followed by detection using 2,2’-azino-bis-3-ethylbenzothiazoline-6-sulfonic acid (ABTS) solution (Pierce, Rockford, IL, USA). Absorbance was measured at 405 nm with a Multiscan Ascent microplate reader (Lab-systems, Helsinki, Finland). As a control, binding activity to BSA (Bovine Serum Albumin) was also tested according to the protocol described above. The expression of intact scFv molecules was confirmed by anti-HA (human influenza hemagglutinin). Two replicate experiments were conducted for all the experimental conditions.

## Acknowledgements

This research was supported by Global Research Development Center Program through the National Research Foundation of Korea (NRF) funded by the Ministry of Science and ICT(MSIT) (2015K1A4A3047345).

## Authors’ contributions

J.N. mainly wrote the paper and participated in the development of the platform. O.K., Y.J., and J.E.K. participated in the development of the platform. H.H. developed and implemented the bioinformatic part of the platform. S.K. did biological experiments for the paper and participated in the writing. S.L., J.P., R.H.J., J.P., and E.K. participated in the experiments for the development of the platform. S.K., J.H., D.K.Y. did the biological experiment for the paper. H.L. analyzed the experimental data. A.C.L. participated in the writing and data analysis. T.R., J.C., and S.K. devised the platform and designed the experiments.

